# Loss of FXR1 and FXR2 promotes accumulation of TDP-43 in aging-related stress granules

**DOI:** 10.1101/2025.10.27.684798

**Authors:** Sonja Menge, Lorena Decker, Karin M. Danzer, Patrick Oeckl, Axel Freischmidt

## Abstract

Cytoplasmic mislocalization and aggregation of the RNA-binding protein TDP-43 in vulnerable neurons may accompany the primary neuropathology of various neurodegenerative diseases, or represent the hallmark of others, such as amyotrophic lateral sclerosis. Aging is the major risk factor for neurodegeneration, and sufficient to induce accumulation of TDP-43 in chronic cytoplasmic stress granules possibly further maturing to irreversible aggregates. Reduced expression of Fragile X protein (FXP) family members (FMR1, FXR1 and FXR2) in vulnerable neurons is associated with neurodegeneration, and loss of each individual FXP induces overlapping and unique aging-related phenotypes in HAP1 and SH-SY5Y cell models. Therefore, we analyzed consequences of FXP loss on TDP-43 cytoplasmic mislocalization and stress granule formation in fibroblast-like HAP1 FXP knockout cells. FXP loss induced nuclear pore pathology, and passive egress of proteins and RNA was evident upon loss of FXR1 or FXR2. Cytoplasmic mislocalization of TDP-43 was restricted to FXR1 knockout cells upon impairment of nuclear import, and cytoplasmic TDP-43 induced spontaneous stress granules exclusively in FXR2 knockout cells. In contrast, loss of FMR1 had no effect on nucleocytoplasmic exchange or TDP-43. Hence, reduced expression of FXR1 and FXR2 in aging and neurodegeneration may contribute to TDP-43 pathology.

## Introduction

Cytoplasmic mislocalization and possible aggregation of Transactive response DNA binding protein of 43 kDa (TDP-43), a predominantly nuclear RNA-binding protein (RBP) involved in most aspects of RNA metabolism and beyond, may accompany the primary neuropathology of various neurodegenerative diseases (NDDs). In amyotrophic lateral sclerosis (ALS), frontotemporal dementia (FTD) with TDP-43- immunoreactive pathology (FTD-TDP), limbic-predominant age–related TDP-43 encephalopathy (LATE) and Perry syndrome, TDP-43 pathology in vulnerable neurons and tissues is the hallmark of the respective disease (Nag and Schneider, 2023). Specific variants in *TARDBP* coding for TDP-43 cause ALS in some rare cases (Kabashi et al., 2008), but in the vast majority (>95%) of patients, wildtype TDP-43 mislocalizes and aggregates in vulnerable neurons (Al-Chalabi et al., 2012; Hardiman et al., 2017).

Similar to other NDDs, aging is the major risk factor for developing ALS, and pathogenic mechanisms overlap with hallmarks of aging at the molecular level. These include, but are not limited to, genetic and epigenetic alterations, loss of proteostasis, mitochondrial dysfunction, impaired autophagy, and cellular senescence (Jagaraj et al., 2024). A very recent study showed that aged neurons express lower level of thousands of RBPs, and nuclear RBPs associated with splicing mislocalize to the cytoplasm. This includes TDP-43 that is incorporated into chronic cytoplasmic stress granules (SGs) that impair proper stress responses, and may represent precursors of TDP-43 aggregates found in ALS and other NDDs (Rhine et al., 2025).

The same study (Rhine et al., 2025) reports that all three members of the Fragile X protein (FXP) family comprising FMR1, FXR1 and FXR2 are among depleted RBPs in aged neurons. The FXPs are predominantly cytoplasmic RBPs regulating local translation at dendrites and synapses, and multiple additional functions have been described (Majumder et al., 2020; Richter and Zhao, 2021; Mueller et al., 2023). These proteins have been repeatedly and independently linked to ALS pathogenesis by different unbiased approaches, i.e. by determining binding partners of ALS-related proteins (Blokhuis et al., 2016) and microRNAs (Freischmidt et al., 2021), and by screening RBPs for modulators of TDP-43 toxicity in *Drosophila* (Coyne et al., 2015). Moreover, overexpression of FMR1 in FUS- and TDP-43-linked zebrafish (Blokhuis et al., 2016) and *Drosophila* (Coyne et al., 2015) ALS models, respectively, substantially rescued respective phenotypes. In ALS post-mortem spinal cord tissue, our group reported aberrant expression of FXR1 and FXR2 in motoneurons independently of the underlying disease gene (Freischmidt et al., 2021), and single-cell proteomics stratified by TDP-43 pathology indicates downregulation of both FXR1 and FXR2 in motoneurons harboring pathogenic aggregates (Guise et al., 2024). Known functions of the FXPs suggest involvement of these proteins in ALS pathogenesis at multiple levels, but in most cases direct evidence is lacking so far (reviewed in (Mueller et al., 2023)).

Recently, we performed an unbiased comparison of downstream events associated with the loss of each individual FXP in non-neuronal HAP1 and neuron-like SH-SY5Y cell lines (Menge et al., 2025). Thereby, we found that regulation of processes linked to cellular senescence and aging may represent another important function of this protein family, and defects induced by the loss of individual FXPs largely overlap with mechanisms generally defective in NDDs including ALS (Gan et al., 2018; Wilson et al., 2023). Specifically, we found that loss of the FXPs is related to defects in ribosome biogenesis, decreased autophagy and impaired lysosome biology, mitochondrial dysfunction, and premature senescence characterized by increased DNA damage, increased expression and/or activity of senescence-associated ß-Galactosidase, and accumulation of lipofuscin. However, analyses of total and insoluble proteomes of HAP1 cells, as well as additional assays, did not indicate any increase in protein aggregation upon loss of individual FXPs. This includes major proteins aggregating in NDDs, such as TDP-43 (Menge et al., 2025). Consequently, in this study, we investigate effects of FXP loss on TDP-43 in more detail.

## Results

### FXP-ko HAP1 cells are similarly depleted in RNA-binding proteins like aging neurons and fibroblasts

Previously, we thoroughly characterized the control (Ctl) and FXP knockout (FXP-ko) HAP1 cells used in this study (Menge et al., 2025). HAP1 is a near-haploid, fibroblast-like cell line derived from chronic myelogenous leukemia cell line KBM-7 that no longer expresses hematopoietic markers (Carette et al., 2011). Importantly, the recently reported aging-induced dysregulation of RBP expression and localization was not restricted to neurons, but was also evident in fibroblasts, although less pronounced (Rhine et al., 2025). Consequently, we first investigated whether premature senescence induced by loss of each individual FXP also leads to a general depletion of RBPs in HAP1 cells. For this, we re-analyzed our proteomic data published previously (Menge et al., 2025), but did not use a fold change cut-off facilitating detection of more subtle changes. The complete analysis of 4656 reliably detected and quantified proteins can be found in Supplementary Table 1. The Gene Ontology (GO) term ‘*RNA binding*’ (GO:0003723) was indeed the most significantly enriched molecular function among downregulated proteins in all three FXP-ko cell lines. RBPs were also enriched in upregulated proteins, but this was substantially less pronounced (Figure 1A). Additionally, we analyzed changes in the expression of RBPs and non-RBPs (according to GO term ‘*RNA binding*’; GO:0003723) in FXP-ko cells compared to controls, and found stronger downregulation of RBPs compared to all other proteins in all three FXP-ko cell lines. RBP depletion was most and least pronounced upon ko of FXR2 and FMR1, respectively, but this difference did not reach statistical significance (Figure 1B). Specifically for TDP-43, proteomics (Figure 1C; Supplementary Table 1) and Western blotting (Figure 1D) did not indicate significant expression changes in FXP-ko HAP1 cells. Taken together, FXP-ko HAP1 cells mirror age-related RBP depletion reported in neurons and fibroblasts (Rhine et al., 2025), and may be suitable for studying aging- and FXP-dependent effects on TDP-43.

**Figure 1.**
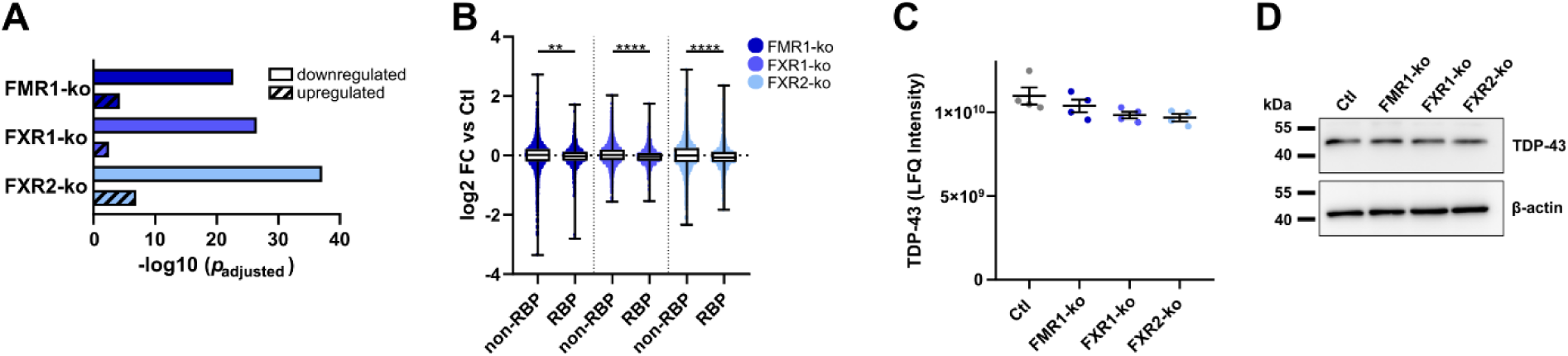
FXP-ko HAP1 cells are globally depleted in RBPs, but expression of TDP-43 is unchanged. (**A**) Enrichment of RBPs (Gene Ontology term ‘*RNA binding*’; GO:0003723) among up- and downregulated proteins in FXP-ko HAP1 cells. (**B**) Expression changes (mean log_2_ fold change from 4 independent experiments) of RBPs (n = 1064) and non-RBPs (n = 3592) in the respective FXP-ko HAP1 cell lines compared to control cells (n = 4; boxes indicate the range of the 25^th^ and 75^th^ percentiles of all data points; the centerline represents the median and whiskers show the full range of all data points; ** *p* < 0.01, **** *p* < 0.0001 in a Kruskal– Wallis test followed by Dunn’s multiple comparisons test). (**C**) Label-free quantification intensities of TDP-43 in FXP-ko cell lines as determined by proteomics (n = 4; bars indicate mean ± s.e.m.; non-significant in a one-way ANOVA). (**D**) Western blot of TDP-43 in FXP-ko HAP1 cell lines.

### Loss of individual FXPs has no effect on the cellular distribution of endogenous TDP- 43 and ALS-related TDP-43 M337V

Next, we addressed subcellular localization and possible incorporation of endogenous TDP-43 into spontaneous cytoplasmic SGs using immunocytochemistry (ICC). However, there was no difference in TDP-43 nucleocytoplasmic distribution or cytoplasmic puncta between FXP-ko and control cells (Figure 2A-C). Hypothesizing that effects of FXP loss on TDP-43 are unmasked when an ALS-associated pathogenic variant is expressed, we overexpressed myc-tagged TDP-43 M337V in the HAP1 cell lines. However, here too, nucleocytoplasmic distribution and cytoplasmic puncta were unchanged in FXP-ko cells compared to controls (Figure 2D-F). From these data we conclude that either premature senescence induced by loss of individual FXPs is not sufficient to induce cytoplasmic mislocalization of splicing proteins, or that fibroblast-like HAP1 cells do not resemble this finding from neurons and primary fibroblasts (Rhine et al., 2025).

**Figure 2.**
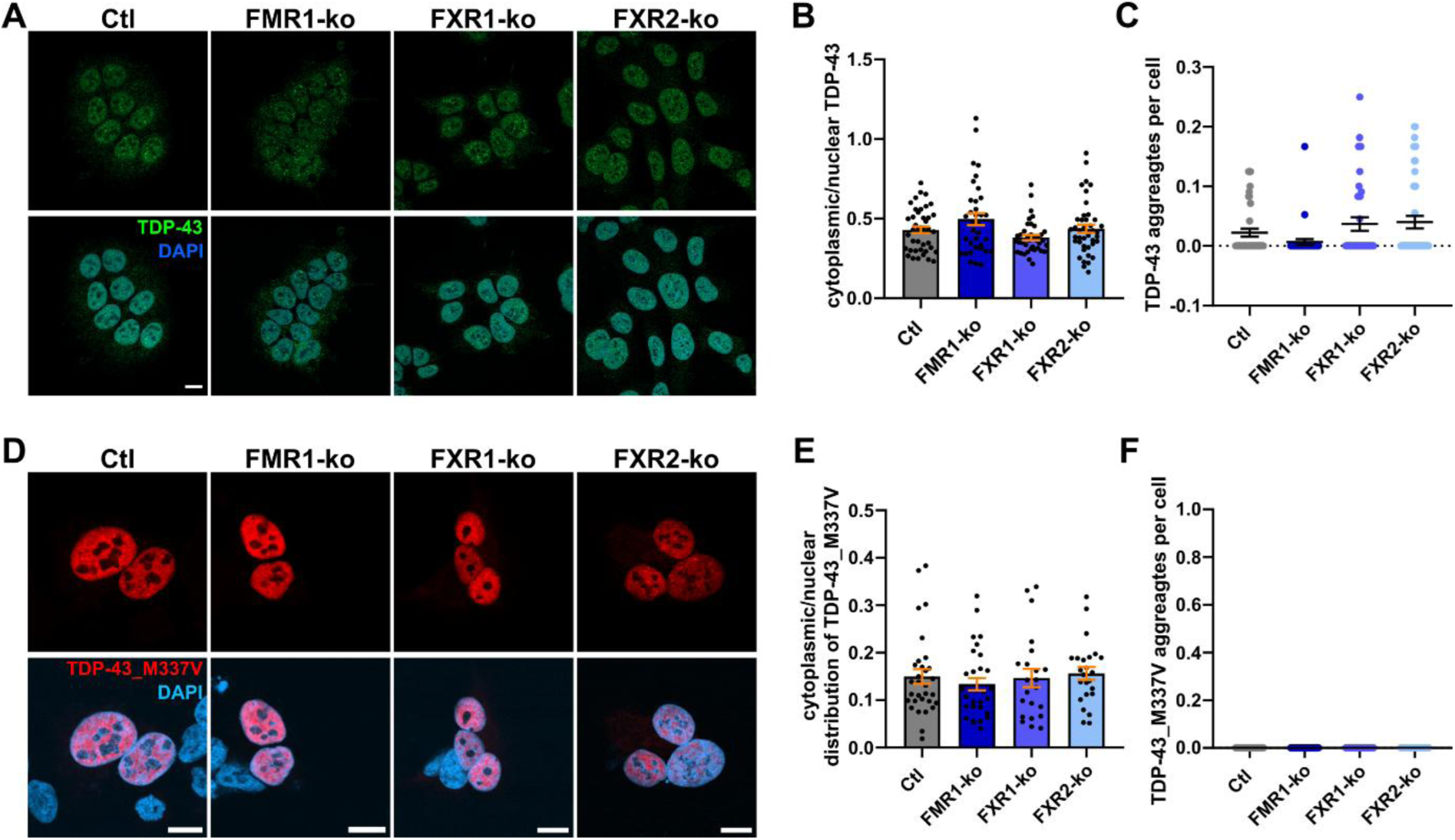
Loss of individual FXPs has no effect on the nucleocytoplasmic distribution and cytoplasmic puncta formation of endogenous TDP-43 and exogenous TDP-43 M337V. (**A-C**) Representative images (**A**) and quantification of nucleocytoplasmic distribution (**B**) and cytoplasmic puncta formation (**C**) of endogenous TDP-43 in respective untreated FXP-ko HAP1 cell lines (n = 28-40 images from 4 independent experiments). (**D-F**) Representative images of respective FXP-ko HAP1 cell lines expressing N-terminally myc-tagged TDP-43 M337V for 24 h (**D**). Nucleocytoplasmic distribution (**E**) and cytoplasmic puncta formation (**F**) of exogenous (anti-myc antibody) TDP-43 M337V are shown (n = 23-35 images from 3-4 independent experiments; bars indicate mean ± s.e.m.; non-significant in a one-way ANOVA; scale bars = 10 µm).

### Cytoplasmic TDP-43 links age-related chronic SGs to the loss of FXR1 and FXR2

TDP-43 is predominantly localized in the nucleus while the FXPs are mostly found in the cytoplasm (Uhlén et al., 2015), and TDP-43 has been shown to interact with all three FXPs (Blokhuis et al., 2016). Additionally, TDP-43 as well as all three FXPs are components of cytoplasmic SGs (Jain et al., 2016). Therefore, we hypothesized that the FXPs may not be involved in the cytoplasmic mislocalization of TDP-43, but in the formation of age-related chronic SGs. To test this, we expressed myc-tagged TDP-43 with a non-functional nuclear localization signal (TDP-43 ΔNLS) (Winton et al., 2008) in the HAP1 cell lines. While all cell lines showed the expected partial redistribution of TDP-43 from the nucleus to the cytoplasm, we unexpectedly detected increased cytoplasmic TDP-43 in FXR1-ko cells (Figure 3A,B). Additionally, in FXR2-ko cells, we found increased numbers of cytoplasmic puncta of TDP-43 that were phosphorylated (Serine 409/410); Figure 3A,C), and reminiscent of aging-induced chronic SGs recently discovered (Rhine et al., 2025). Colocalization of cytoplasmic TDP-43 accumulations with SG marker TIAR suggests that loss of FXR2, but not of FMR1 or FXR1, leads to SG formation in the absence of exogenous stressors (Figure 3D-F). Interestingly, increased SG formation was not detectable in FXR2-ko cells without expression of cytoplasmic TDP-43 (Figure 3G,H). Hence, cytoplasmic TDP-43 may induce formation of SGs in FXR2-ko cells.

**Figure 3.**
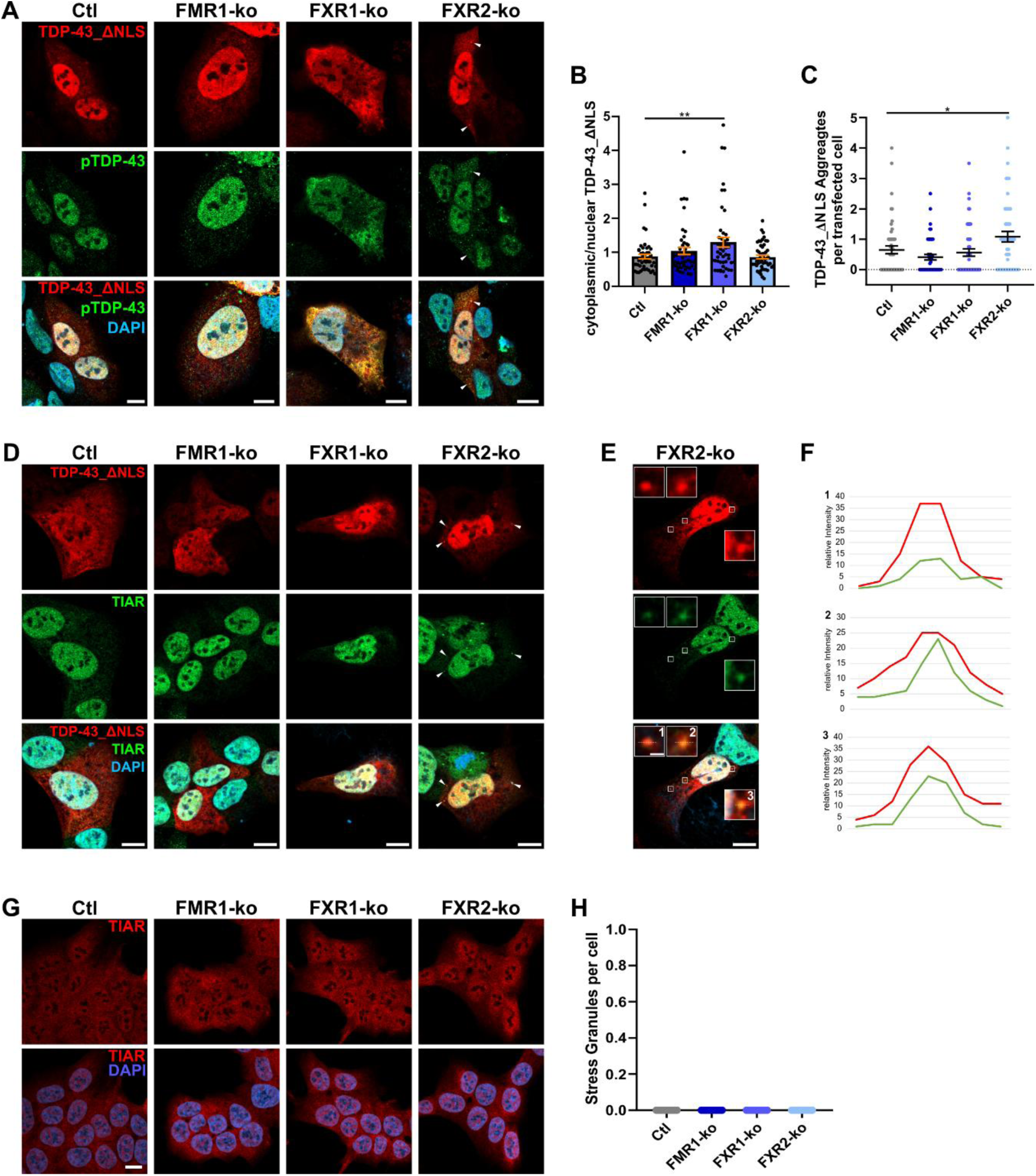
TDP-43 ΔNLS shows increased cytoplasmic localization in FXR1-ko, and induces stress granule formation in FXR2-ko HAP1 cells. (**A-C**) Representative images of respective FXP-ko HAP1 cell lines expressing N-terminally myc-tagged TDP-43 ΔNLS for 24 h. Staining with anti-myc and anti-phosphoTDP-43 (Serine 409/410) antibodies are shown (**A**). Quantification of nucleocytoplasmic distribution (**B**) and cytoplasmic puncta formation (**C**) of exogenous TDP-43 ΔNLS (n = 45 images from 5 independent experiments). (**D-F**) Representative images of respective HAP1 cell lines expressing myc-tagged TDP-43 ΔNLS for 24 h. Co-staining with anti-myc and anti-TIAR antibodies are shown (**D**). Magnification (**E**) and intensity analyses (**F**) of anti-myc and anti-TIAR signals in FXR2-ko cells. (**G, H**) Representative images (**G**) and numbers of stress granules per cell (**H**) of respective untreated FXP-ko HAP1 cells stained with anti-TIAR antibody (n = 50 images from 5 independent experiments; bars indicate mean ± s.e.m.; \**p* < 0.05, \*\**p* < 0.01 in a one-way ANOVA followed by *post hoc* Šídák’s test; scale bars = 10 µm).

### Sorbitol induced stress replicates findings from TDP-43 ΔNLS

Next, we aimed at validating these results at the endogenous level, and used osmotic stressor sorbitol to induce cytoplasmic SGs. Similar to other cell lines (Dewey et al., 2011), sorbitol robustly induced TDP-43 containing SGs also in HAP1 cells. Quantification of both SG numbers and nucleocytoplasmic distribution of TDP-43 replicated findings from overexpressing TDP-43 ΔNLS, i.e. we found increased cytoplasmic TDP-43 in FXR1-ko cells (Figure 4A,B), and increased numbers of SGs in FXR2-ko cells (Figure 4A,C). In FMR1-ko cells, nucleocytoplasmic distribution of TDP- 43 was comparable to the controls, but SG numbers were decreased. Despite the differences in SG numbers and localization of TDP-43, Western blotting of soluble and insoluble TDP-43 under basal and stressed conditions did not reveal any differences between the cell lines. While under basal conditions the vast majority of TDP-43 was soluble, sorbitol treatment induced insolubility of virtually all TDP-43 (Figure 4D,E).

**Figure 4.**
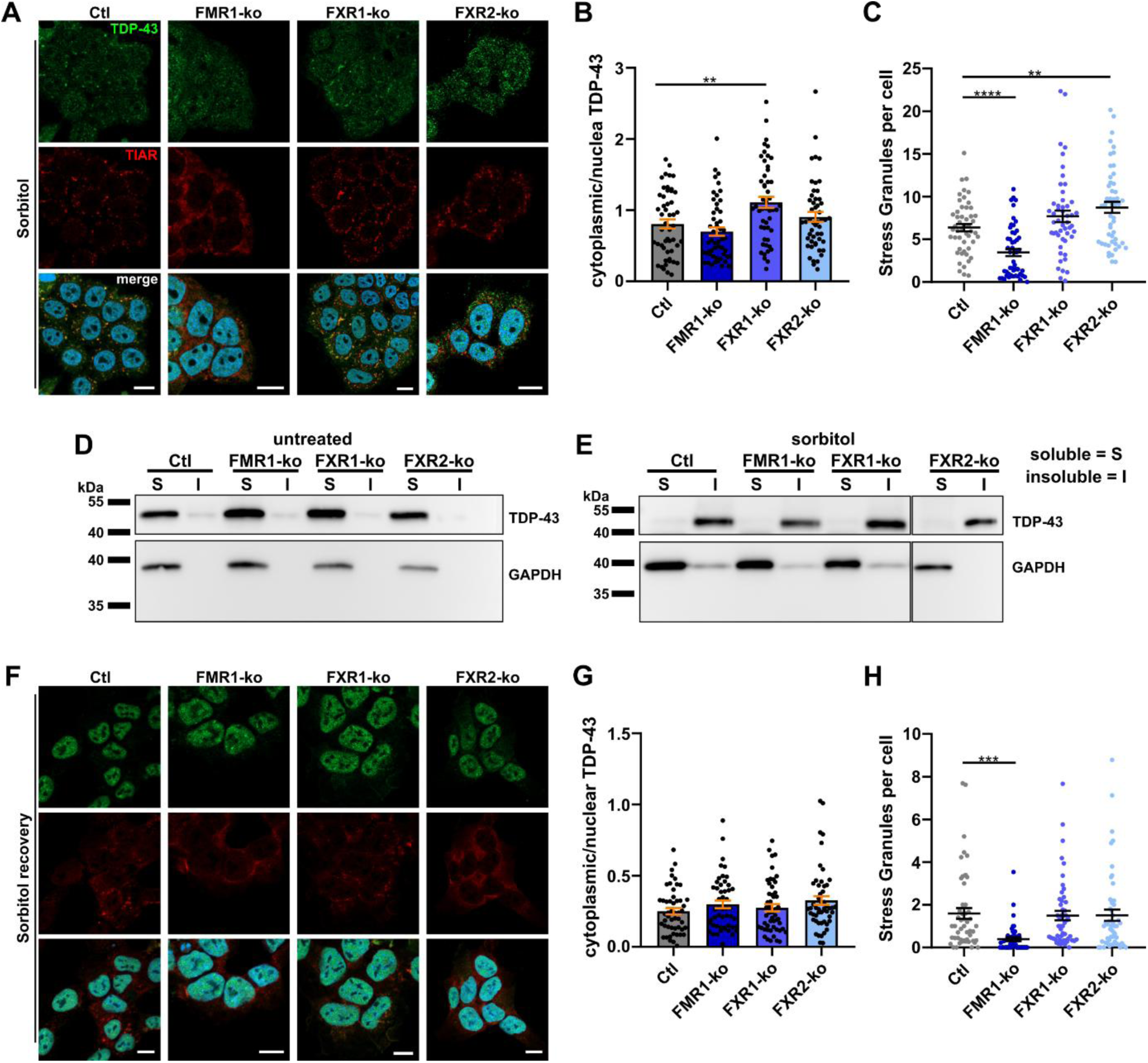
Stress granule induction using sorbitol leads to increased cytoplasmic TDP-43 in FXR1-ko, and increased numbers of stress granules in FXR2-ko HAP1 cells. (**A-C**) Representative images of respective HAP1 cell lines treated with sorbitol (400 mM for 2 h; **A**). Quantification of TDP-43 nucleocytoplasmic distribution (**B**) and numbers of stress granules per cell (**C**) are shown (n = 50 images from 5 independent experiments). (**D, E**) Western blots of RIPA soluble (S) and insoluble (I) fractions of endogenous TDP-43 in untreated (**D**) and sorbitol treated (400 mM for 2 h; **E;** the vertical line indicates that data from FXR2-ko cells is from a different blot) HAP1 cell lines. GAPDH is shown as a marker for the soluble fraction. (**F-H**) Representative images of HAP1 cells 30 min after release of sorbitol stress (400 mM for 2 h; **F**). Quantification of TDP-43 nucleocytoplasmic distribution (**G**) and numbers of stress granules per cell (**H**) are shown (n = 50 images from 5 independent experiments; bars indicate mean ± s.e.m.; \*\**p* < 0.01, \*\*\**p* < 0.001, \*\*\*\**p* < 0.0001 in a one-way ANOVA followed by *post hoc* Šídák’s test; scale bars = 10 µm).

To reveal possible impairments in nuclear re-localization of TDP-43 and in SG clearance, we also quantified nucleocytoplasmic distribution of TDP-43 and SG numbers upon release of sorbitol stress. As early as 30 min after stress release, differences between the cell lines were no more detectable, except the lower numbers of SGs in FMR1-ko cells (Figure 4F-H). From these data we conclude that SG clearance is not affected by FXP loss, and that active nuclear import of TDP-43 is still functional in FXR1-ko cells. Hence, in FXR1-ko cells, other mechanisms may lead to increased cytoplasmic TDP-43 when nuclear import is decreased, e.g. by deleting the NLS, or when cellular stress leads to cytoplasmic redistribution of RBPs.

### Loss of individual FXPs induces nuclear envelope pathology

Next, we addressed mechanisms that may be responsible for increased cytoplasmic TDP-43 in FXR1-ko cells when nuclear import is impaired. All three FXPs, and especially FXR1, have recently been shown to regulate nucleoporin localization and nuclear envelope morphology during interphase of the cell cycle (Agote-Aran et al., 2020). Here, knockdown of FXPs led to irregularly shaped nuclei and formation of cytoplasmic nucleoporin condensates. These changes in nuclear envelope architecture had no effect on steady-state nucleocytoplasmic transport similar to our observations of TDP-43 localization in FXR1-ko cells described above. Consequently, we investigated nuclear envelopes and possible formation of nucleoporin condensates in the FXP-ko cell lines using MAb414 antibody detecting a range of nucleoporins containing phenylalanine-glycine (FG) repeats (FG-nups). Surprisingly, in contrast to the previous report (Agote-Aran et al., 2020), we found decreased instead of increased numbers of cells containing FG-nup condensates in all FXP-ko cell lines (Figure 5A,B). Nevertheless, loss of each FXP induced nuclear envelope pathology assessed by counting cells with a blurry appearance or interruptions of MAb414 stained nuclear envelopes (Figure 5C,D). Additionally, nuclei of FMR1-ko cells were less round compared to controls (Figure 5E).

**Figure 5.**
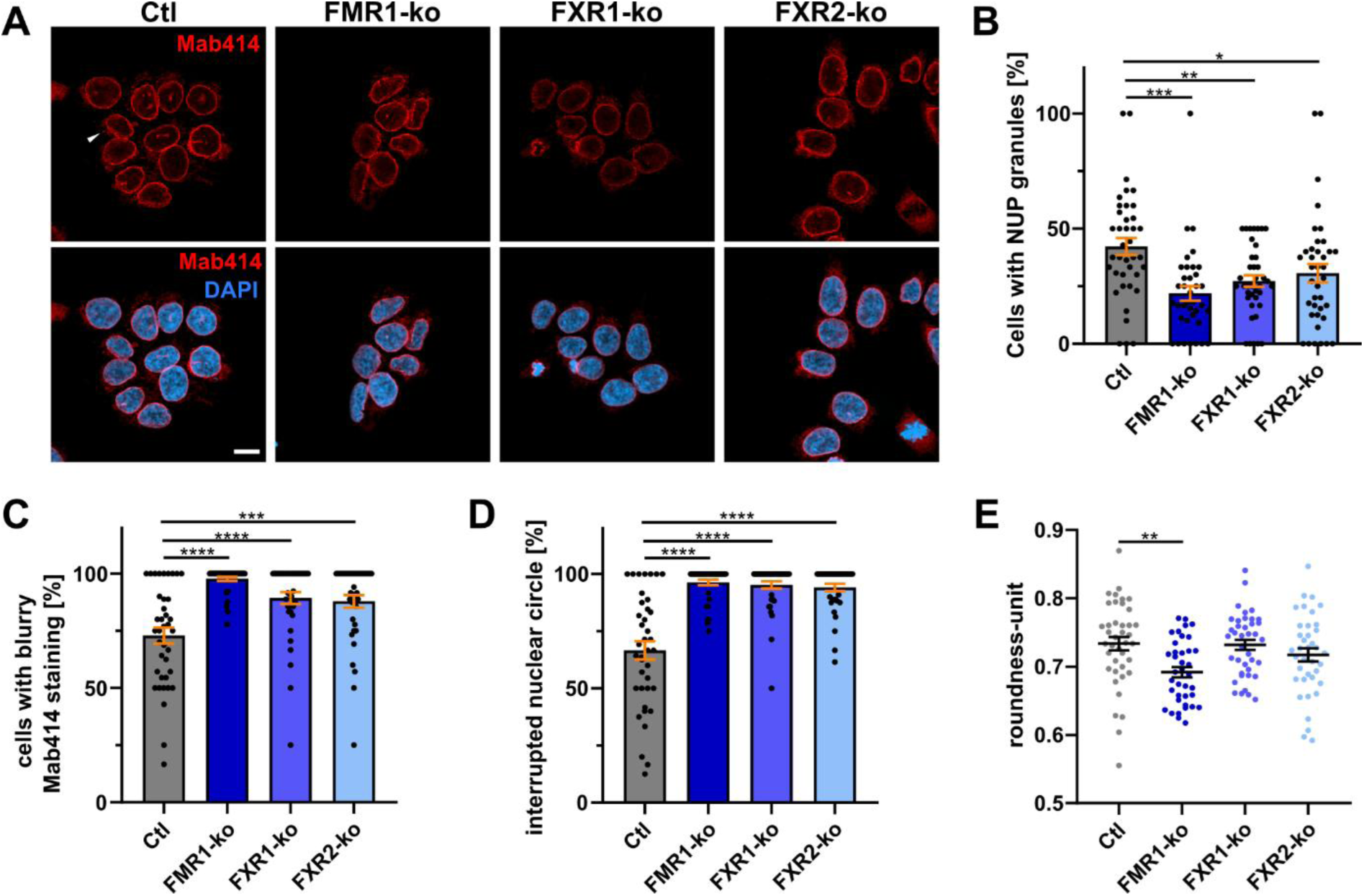
Loss of individual FXPs induces nuclear envelope pathology in HAP1 cells. Representative images of respective HAP1 cell lines stained with Mab414 antibody (**A**). Quantification of cells showing cytoplasmic accumulations of phenylalanine-glycine repeat containing nucleoporins (**B**), blurry appearance (**C**) or interruptions of nuclear envelopes (**D**). (**E**) Determination of roundness of nuclei (n = 38-40 images from 4 independent experiments; bars indicate mean ± s.e.m.; \**p* < 0.05, \*\**p* < 0.01, \*\*\**p* < 0.001, \*\*\*\**p* < 0.0001 in a one-way ANOVA followed by *post hoc* Šídák’s test; scale bar = 10 µm).

### Loss of FXR1 and FXR2 enhances passive egress of proteins from the nucleus

FG-nups form a complex meshwork in the central channel of nuclear pores providing a permeability barrier for nucleocytoplasmic exchange. Generally, smaller proteins (< ≈40 kDa) can cross this barrier by passive diffusion while larger proteins require nuclear transport receptors (Li et al., 2016). To test if changes in passive diffusion may explain increased cytoplasmic TDP-43 upon loss of nuclear import, we expressed EGFP (≈27 kDa) that does not contain a NLS in the FXP-ko cell lines and quantified nucleocytoplasmic distribution. Here, increased cytoplasmic EGFP was detected in FXR1- and FXR2-ko cells (Figure 6A,B). We also determined nucleocytoplasmic distribution of endogenous p62 (*SQSTM1*) that does not contain a NLS and shows predominant cytoplasmic localization. Here too, similar to EGFP, p62 was more cytoplasmic in FXR1- and FXR2-ko cells indicating that passive exchange is enhanced exclusively out of the nucleus and not following concentration gradients (Figure 6C,D). Hence, more “leaky” nuclear pores may contribute to increased cytoplasmic TDP-43 in FXR1-ko cells upon abrogation of active transport. Although FXR2-ko cells showed similar defects in nucleocytoplasmic exchange, enhanced passive egress alone cannot account for the increased cytoplasmic TDP-43 observed specifically in FXR1-ko cells.

**Figure 6.**
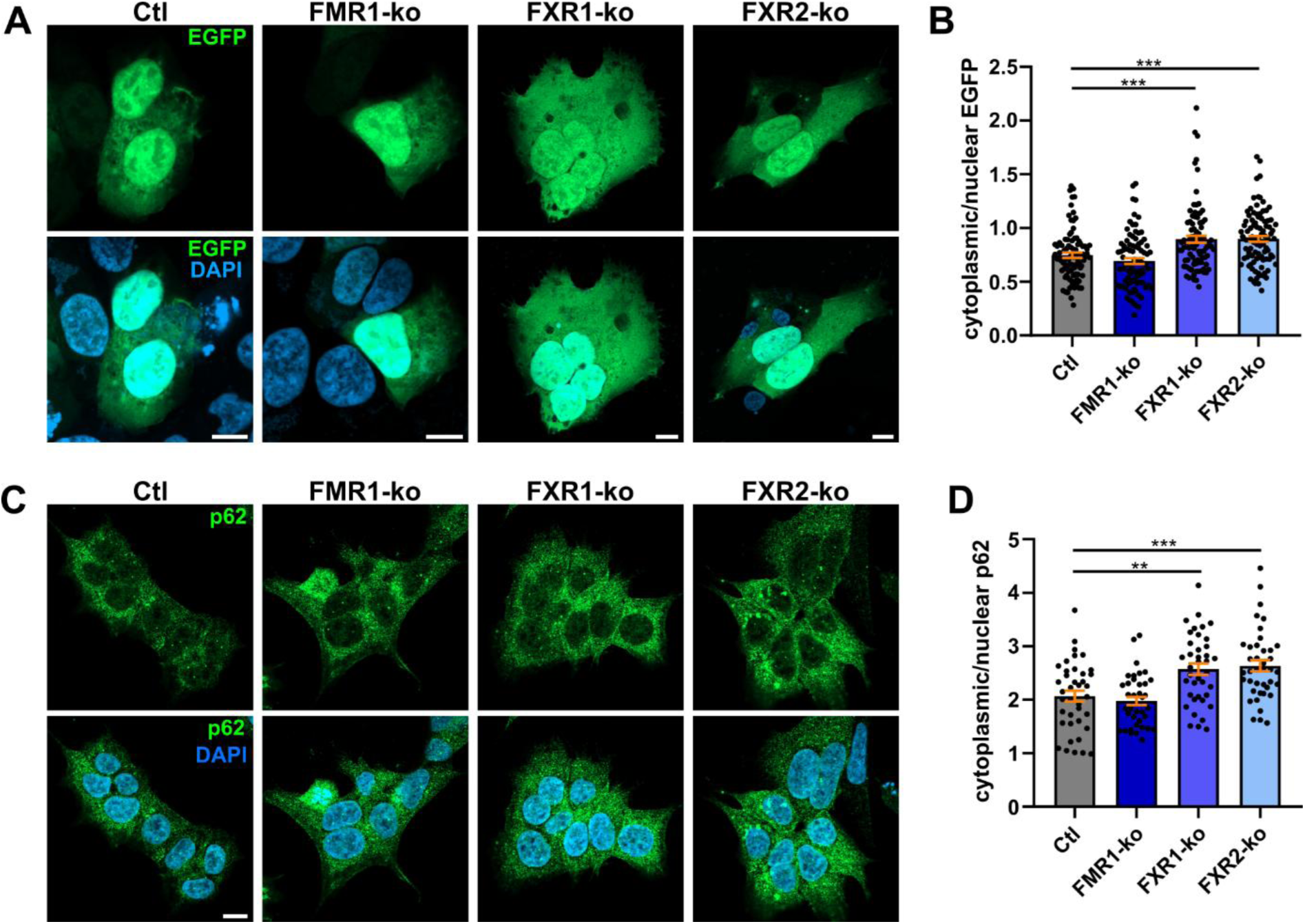
Loss of FXR1 or FXR2 leads to more leaky nuclei. (**A, B**) Representative images of HAP1 cell lines expressing EGFP for 24 h (**A**), and respective nucleocytoplasmic distribution of EGFP (**B**; n = 90 images from 9 independent experiments). (**C, D**) Representative images (**C**) and nucleocytoplasmic distribution (**D**) of endogenous p62 (*SQSTM1*) in untreated FXP- ko HAP1 cell lines (n = 40 images from 4 independent experiments; bars indicate mean ± s.e.m.; \*\**p* < 0.01, \*\*\**p* < 0.001 in a one-way ANOVA followed by *post hoc* Šídák’s test; scale bar = 10 µm).

### Loss of FXPs decreases transcription

Besides active nuclear import, nuclear localization of TDP-43 is also partly dependent on binding to nuclear RNAs, and loss of RNA binding capabilities of TDP-43 or reduced nuclear RNA abundance (Duan et al., 2022) or transcription (Ederle et al., 2018) results in passive nuclear efflux of TDP-43. Aging leads to increased speed of transcription, but total output is reduced (Papantonis et al., 2024). Considering that FXP loss induces some hallmarks of senescence (Menge et al., 2025), we hypothesized that reduced transcription may contribute to increased cytoplasmic TDP-43 in FXR1-ko cells. For testing, we used 5-ethynyluridin (EU) labelling of nascent RNA followed by covalent coupling of EU to a fluorescent dye and detection by microscopy. Here, we observed a reduction in transcriptional output in all FXP-ko cells, but most pronounced in FXR1- and FXR2-ko cells. Results of total and specifically nuclear nascent RNA were very similar (Figure 7A,B,C). Interestingly, by determining nucleocytoplasmic distribution, we found increased cytoplasmic localization of nascent RNA in FXR1- and FXR2-ko cells (Figure 7D). Thus, enhanced passive egress from the nucleus is not restricted to proteins, and reduced transcription may contribute to increased cytoplasmic TDP-43. However, again, FXR1- and FXR2-ko cells showed highly similar results.

**Figure 7.**
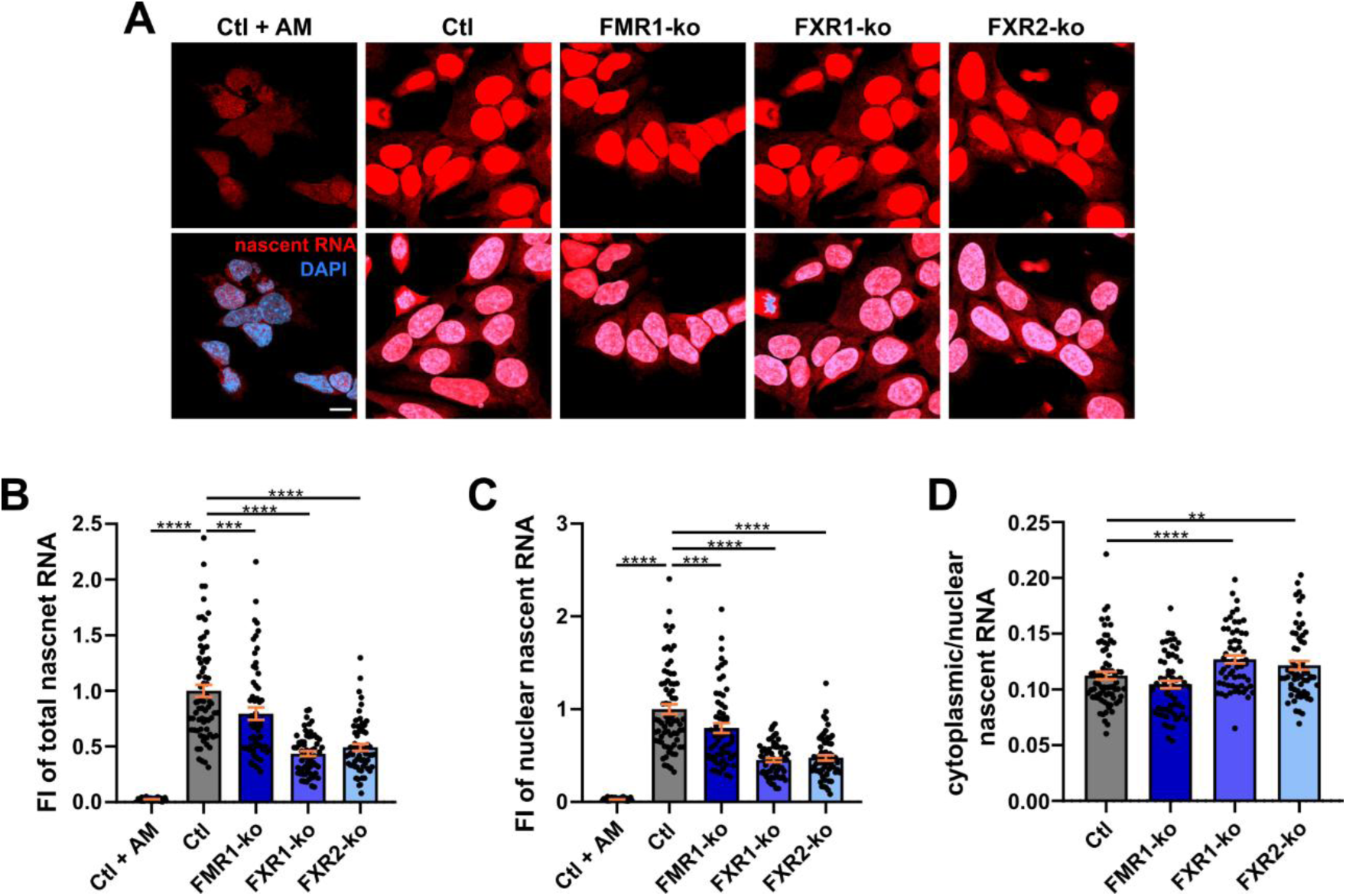
Loss of individual FXPs decreases transcription. (**A**) Representative images of untreated HAP1 cells where nascent RNA was labelled with 5-ethynyluridin followed by coupling of 5-ethynyluridin to a fluorescent dye. Please note that images in this experiment were not overexposed, but are shown here with increased brightness to also visualize nascent RNA in the cytoplasm. Quantification of total (**B**) and nuclear (**C**) nascent RNA, as well as nucleocytoplasmic distribution of nascent RNA (**D**) are shown. Control HAP1 cells treated with transcription inhibitor Actinomycin-D (AM; 100 ng/ml for 18 h) are shown as control (n = 40 images from 4 independent experiments; bars indicate mean ± s.e.m.; \*\*\**p* < 0.001, \*\*\*\**p* < 0.0001 in a one-way ANOVA followed by *post hoc* Šídák’s test; scale bar = 10 µm).

### Proteomics does not explain differences in TDP-43 nucleocytoplasmic distribution and SG formation in FXP-ko cells

Despite more “leaky” nuclear pores and decreased transcription in respective FXP-ko cells, this does not explain why increased nuclear efflux of TDP-43 is restricted to the loss of FXR1, and why chronic SGs form exclusively in FXR2-ko cells. Therefore, we analyzed expression of proteins involved in nuclear pores and nucleocytoplasmic transport (KEGG pathway “*nucleocytoplasmic transport*”; hsa03013), as well as components of SGs (GO term “*cytoplasmic stress granule*”; GO:0010494) in our proteomic data. 85 out of 108 (79%) proteins involved in nucleocytoplasmic transport, and 66 out of 150 (44%) SG components were reliably detected and quantified in all three FXP-ko HAP1 cells.

PCA of proteins involved in nucleocytoplasmic transport surprisingly revealed that changes are most pronounced in FMR1- and FXR2-ko cells, while FXR1-ko cells partly overlap with FXR2-ko and control cells (Figure 8A). At the level of individual proteins, not a single one was exclusively dysregulated in FXR1-ko cells. Without applying multiple testing correction, the nucleoporin SEH1L was slightly downregulated in FXR1-ko cells only. However, fold changes indicated a comparable decrease of SEH1L also in FMR1- and FXR2-ko cells, and this protein is unlikely to account for increased cytoplasmic TDP-43 exclusively in FXR1-ko cells (Figure 8B).

**Figure 8.**
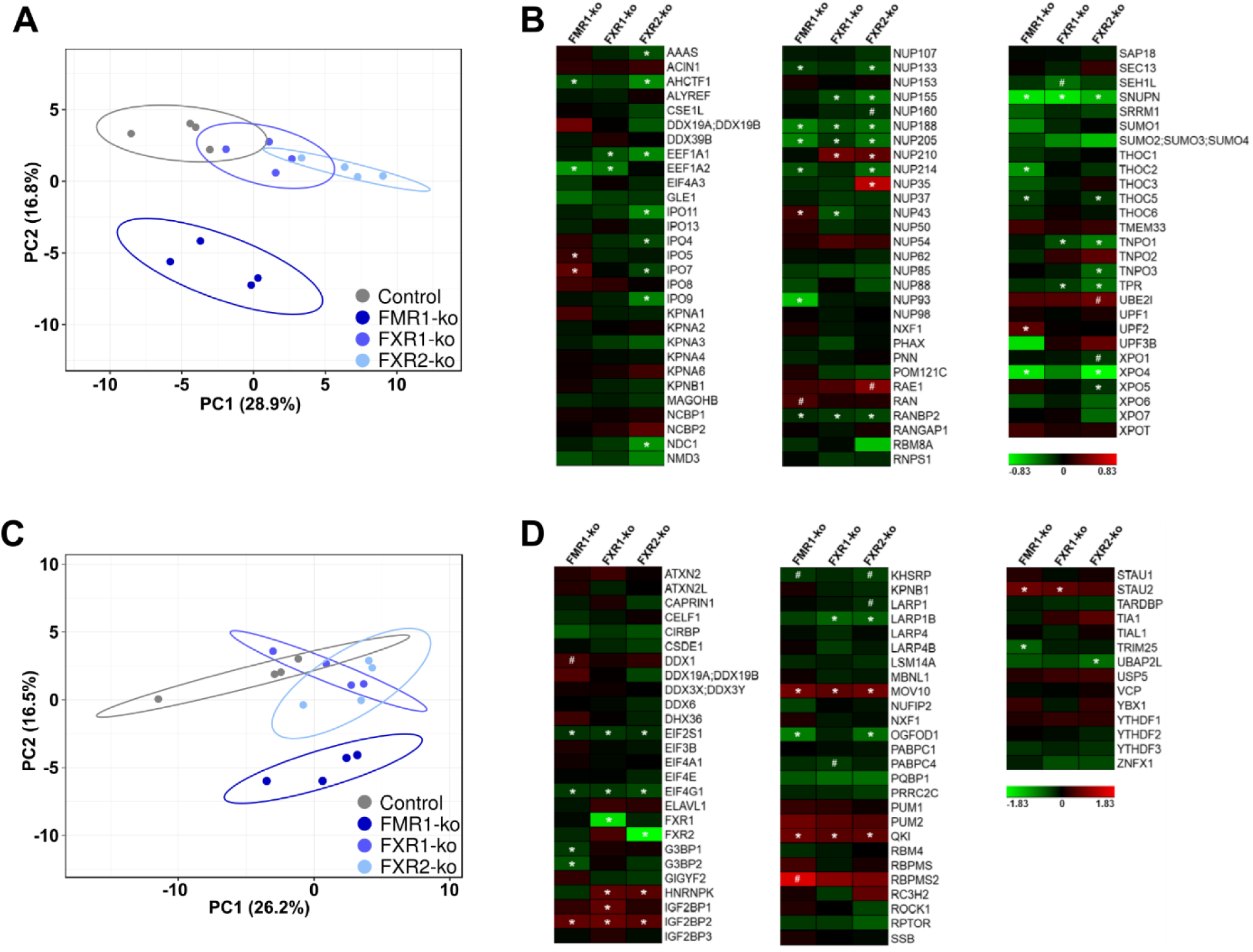
Separate analyses of proteins involved in nucleocytoplasmic transport and of stress granule components from proteomic data. (**A, B**) Principal component analysis (**A**) and expression changes compared to control cells (**B**) of proteins involved in nucleocytoplasmic transport (KEGG pathway “*nucleocytoplasmic transport*”; hsa03013; 85 out of 108 reliably detected and quantified). (**C, D**) Principal component analysis (**C**) and expression changes compared to control cells (**D**) of stress granule components (Gene Ontology term “*cytoplasmic stress granule*”; GO:0010494); 66 out of 150 reliably detected and quantified). Color coding of heatmaps in (**B**) and (**D**) represents mean log_2_ fold changes compared to control cells from 4 independent experiments. Statistically significant changes with (*) and without (^#^) adjusting ANOVA *p*-values for multiple testing are indicated.

Regarding components of cytoplasmic SGs, PCA indicated relatively little changes in FXR1- and FXR2-ko cells compared to controls. Here too, most severe changes were evident in FMR1-ko cells (Figure 8C). The only proteins exclusively dysregulated in FXR2-ko cells were UBAP2L and, when disregarding correction for multiple testing, LARP1 (Figure 8D). However, there was a strong trend towards downregulation of UBAP2L in FMR1- and FXR1-ko cells, too, and downregulation of LARP1 was minimal (log2 fold change = -0.175). Furthermore, UBAP2L is essential for the nucleation of SGs (Asano-Inami et al., 2023), and downregulation can thus not explain increased numbers of SGs in FXR2-ko cells. Hence, our proteomic data could not identify promising candidates explaining differences in nucleocytoplasmic exchange and SG formation in FXR1- and/or FXR2-ko cells, respectively.

## Discussion

In this study, we explored consequences of FXP loss on cytoplasmic mislocalization and accumulation of TDP-43 in SGs. While ko of FMR1 had little to no effect on TDP- 43, loss of FXR1 induced enhanced passive nuclear egress of TDP-43, and cytoplasmic TDP-43 induced SGs exclusively in FXR2-ko cells.

In our previous study, we showed that loss of each individual FXP is sufficient to induce premature senescence and other hallmarks of aging in HAP1 cells (Menge et al., 2025). However, aging-associated changes in the different FXP-ko cells were not identical, but loss of each individual FXP induced very specific phenotypes. As FXP loss and aging phenotypes are intrinsically tied to each other in HAP1 cells, we currently cannot differentiate if effects on TDP-43 are due to aging-associated mechanisms, or linked specifically to the loss of the respective FXP. Nevertheless, at least defects exclusively induced by the loss of FMR1, or by all three FXPs, may be excluded to contribute to TDP-43 mislocalization and SG induction. This includes decreased mitochondrial membrane potentials and accumulation of damaged mitochondria, defects in endo-lysosomal pathways and lipofuscin accumulation (FMR1-ko only), as well as increased DNA damage and decreased autophagy (all three FXP-ko cell lines) (Menge et al., 2025). *Vice versa*, TDP-43 mislocalization and accumulation in SGs in aging neurons was accompanied by downregulation of all three FXPs (Rhine et al., 2025) making it difficult to draw conclusions about the contribution of FXP- and/or aging associated mechanisms.

Passive nuclear egress is likely the dominating mechanism for cytoplasmic mislocalization of TDP-43 (Ederle et al., 2018; Pinarbasi et al., 2018), and has been linked to nuclear pore pathology (Coyne et al., 2021; Ramírez-Núñez et al., 2025) as well as decreased nuclear RNA abundance (Duan et al., 2022) and/or transcription (Ederle et al., 2018). However, despite aberrant nuclear envelopes and reduced RNA synthesis in all three FXP-ko cell lines, increased passive nuclear efflux of TDP-43 was restricted to the loss of FXR1. Our proteomic data was not indicative of changes specifically in FXR1-ko cells, but increased reactive oxygen species (ROS) exclusively detected in FXR1-ko cells (Menge et al., 2025) may further drive cytoplasmic mislocalization of TDP-43 (Colombrita et al., 2009; Yan et al., 2025). However, there are various additional explanations for this observation, such as failure of incorporation of TDP-43 in macromolecular complexes preventing its nuclear egress (dos Passos et al., 2024), or loss of FXR1-mediated assembly of mRNPs at nuclear pores (Yang et al., 2025) that may have effects on TDP-43 localization and/or nuclear pore permeability. Importantly, nuclear egress of TDP-43 upon knockout of FXR1 was fully compensated in HAP1 cells, but might become highly relevant in aging and/or neurodegeneration when active nuclear protein import declines (Pujol et al., 2002; Khalil et al., 2024).

SGs have been suspected to be precursors of pathogenic TDP-43 aggregates (Colombrita et al., 2009; Li et al., 2013), and more recent studies still support this hypothesis (Uechi et al., 2025; Yan et al., 2025). The newest discovery of chronic SGs in aging neurons (Rhine et al., 2025) may therefore explain why aging is required for developing NDDs associated with TDP-43 pathology. Here, accumulation of cytoplasmic TDP-43 in SGs was restricted to the loss of FXR2, and cytoplasmic TDP- 43 even induced SG formation although TDP-43 is not known as a SG nucleating protein (Colombrita et al., 2009). In contrast, decreased SG numbers in FMR1-ko cells can be explained by SG nucleating properties of FMR1 (Mazroui et al., 2002; Didiot et al., 2009). All three FXPs and TDP-43 bind to each other and are components of SGs (Zhang et al., 1995; Blokhuis et al., 2016; Jain et al., 2016). Hence, SG formation in FXR2-ko cells may be modified directly by differential protein-protein and/or protein-RNA interactions. Alternatively, loss of FXR2 may indirectly promote SG formation, either by dysregulation of important SG components not detected in our proteomic approach, by inducing different post translational modifications of TDP-43 that promote accumulation in SGs (Cohen et al., 2015), or by causing low level of chronic stress decreasing thresholds for SG assembly. Defects in ribosome biogenesis and decreased translational fidelity were most pronounced or exclusively detected, respectively, in FXR2-ko cells (Menge et al., 2025), and may contribute to increased SG formation.

Taken together, our study indicates that aging-associated TDP-43 mislocalization and accumulation in chronic cytoplasmic SGs are distinct processes, and can be assigned to the loss of FXR1 and FXR2, respectively. Considering that loss of these proteins accompanies both aging (Rhine et al., 2025) and NDDs such as ALS (Freischmidt et al., 2021; Guise et al., 2024), it is plausible to hypothesize that maintaining expression of FXR1 and FXR2 in age-related TDP-43 proteinopathies may benefit patients by mitigating TDP-43 pathology and senescence-associated cellular decline.

## Materials and methods

### Proteomic data analyses

Proteomic data from control and FMR1-, FXR1- and FXR2-ko HAP1 cells analyzed in this study were previously published (Menge et al., 2025). Similar to our previous study, we restricted all analyses to 4656 proteins reliably detected in at least 50% (two out of four) of samples from each cell line. For the identification of differentially expressed proteins, we used a one-way ANOVA with *p*-value adjustment for multiple testing (Benjamini and Hochberg, 1995). *Post hoc* Tukey HSD was then used to identify groups with significant changes, and to correct *p*-values for multiple comparisons. To compare expression of RBPs and non-RBPs in the different HAP1 cell lines, the Kruskal-Wallis-Test followed by *post hoc* Dunn’s test was used. For GO enrichment analyses of up- and downregulated proteins, we used ShinyGO 0.82 (Ge et al., 2020). Principal component analyses were performed with ClustVis using standard settings (Metsalu and Vilo, 2015). Heatmaps were generated with Genesis 1.8.1 (Sturn et al., 2002).

### Cell lines and cell culture

CRISPR/Cas9-edited HAP1 cell lines carrying knockouts of FMR1, FXR1 or FXR2, respectively, were obtained from Horizon Discovery (Waterbeach, UK; Supplementary Table 2). HAP1 cells were grown in IMDM supplemented with 10% fetal calf serum under standard conditions. A thorough characterization of these cells was recently published (Menge et al., 2025). Treatments of cells are described in the respective Figure legends.

### Transfection and plasmids

For transfection of HAP1 cells, we used calcium phosphate precipitation as previously described (Jordan and Wurm, 2004) with minor modifications. For expression of N-terminally myc-tagged human TDP-43 M337V and TDP-43 ΔNLS or EGFP, pcDNA3 (Invitrogen; #A-150228) or pCI-neo (Promega; #E1841) were used, respectively. All plasmids were verified by Sanger sequencing.

### Immunocytochemistry (ICC), microscopy and image analyses

ICC of HAP1 cells was performed exactly as described previously (Freischmidt et al., 2021). For confocal microscopy we used the laser-scanning microscope ZEISS LSM 980 with Airyscan 2 (Axio Observer.Z1 / 7) with a Plan-Apochromat 100x/1.40 Oil M27 objective and the Zen blue software (version 3.3).

For quantification of fluorescence intensities, counting of granules/aggregates, and determining roundness/circularity of nuclei we used Fiji (2.14.0 release, 32 bit, 2017 May 30) (Schindelin et al., 2012). Since HAP1 cells predominantly grow in clusters hindering unbiased selection of single cells by Fiji, image analyses were performed at the level of whole images that did not overlap when acquired from the same slide. Each datapoint in the respective graphs represents the mean value of ≈5-20 quantifiable (i.e. complete) cells from the same image (Menge et al., 2025).

### Fractionation of soluble and insoluble proteins

HAP1 cells were washed thoroughly with PBS and lysed in RIPA buffer (50 mM Tris, 150 mM sodium chloride, 0.25% sodium deoxycholate, 1% NP-40, 1 mM EDTA, pH 7.4). Lysates were briefly sonicated and adjusted to a protein concentration of 1 mg/ml.

Next, 200 µl of lysates were centrifuged at 100,000x *g* and 4° C for 30 min, and resulting supernatants represent the soluble fraction. Pellets of insoluble protein were washed with RIPA buffer and centrifuged again at 100.000x *g* and 4° C for 30 min. Protein pellets were then resuspended in 200 µl 8 M urea buffer (20 mM Tris, 8 M urea, pH 8.0), and represent the insoluble fraction. Equal volumes (20 µl) of soluble and insoluble proteins were used for Western blotting.

### Visualization of nascent RNA

For visualization of nascent RNA, we used the Click-iT™ RNA Alexa Fluor 594™ Imaging-Kit (Thermo Fisher Scientific; #C10330) that is based on incorporation of 5- ethynyluridin (EU) in nascent RNA followed by covalent coupling of 5-EU to a fluorescent dye by Click-iT chemistry. Modifications to the manufacturer’s protocol were increased incubation time of cells in 5-EU containing media (from 1 h to 3 h), and prolonged incubation of fixed cells in the Click-iT reaction cocktail (from 30 min to 1 h).

### Antibodies

All antibodies used in this study for ICC and Western blotting are listed in Supplementary Table 3, including manufacturer, catalogue number and working concentration used.

### Statistical analyses

Statistical analyses of proteomic data are described above. For all other experiments, a one-way ANOVA followed by *post hoc* Šídák’s test was used to detect differences between the four HAP1 cell lines and to correct *p*-values for multiple comparisons. *P*- values < 0.05 were considered statistically significant. Statistical analyses and generation of graphs was performed with Graphpad Prism software (version 8.4.3).

## Acknowledgements

We would like to thank Nadine Dreher and Stephen Meier (Department of Neurology, Ulm University, Germany) for excellent technical assistance.

## Funding

This work was supported by the German Research Foundation (DFG; #521487152 to AF; #450627322 and #251293561 to KMD), the TargetALS Foundation (to KMD), the ALS Association/ALS Finding A Cure (#24-SGP-691 and #23-PPG-674-2 to PO), the Charcot Foundation (#D.7090 to PO), the Cure Alzheimeŕs Fund (to PO), the DZNE Innovation-to-Application (#I2A_call9_Oeckl to PO) and the EU Horizon Europe (#SYNAPSING to PO).

## Notes

### Competing Interest Statement

The authors have declared no competing interest.

